# Compartmentalization of Transcripts During Antibody Mediated Rejection in Renal Transplants

**DOI:** 10.1101/2024.05.20.594331

**Authors:** Dajana Margeta, Hirotsugu Noguchi, Sepideh Khazaie, Leal C. Herlitz, Joshua J. Augustine, Peter S. Heeger, Anat R. Tambur, Robert L. Fairchild, William M. Baldwin

## Abstract

We used Digital Spatial Profiling to localize transcripts in glomeruli and tubulointerstitial compartments in a series of 4 biopsies from a patient diagnosed with acute antibody-mediated rejection (AMR). The 4 biopsies included: a baseline protocol biopsy 25 days after transplantation; a 3 month biopsy diagnosed as acute AMR; a biopsy 4 months after treatment with intravenous immunoglobulin (IVIg) showing ongoing AMR with a mild increase in tubulointerstitial fibrosis; and a biopsy 7 months later with resolution of glomerulitis. Glomeruli were captured in regions of interest (ROIs) for whole exome sequencing. Compared to baseline glomeruli, 17 transcripts were increased and 39 decreased in the 3 subsequent biopsies (> 2-fold and p < 0.005). Increased signatures for macrophages correlated with increased numbers of CD68 positive cells imaged in the corresponding glomeruli. The Human Cell Atlas classified the 39 transcripts decreased during the initial rejection as characteristic of podocytes and this gene signature did not recover in the subsequent 2 biopsies. Additional ROIs encompassing areas of tubulointerstitial fibrosis disclosed signatures for memory B cells in the acute AMR sample. Treatment with IVIg did not eliminate the B cell signal in the subsequent biopsy.

Collectively these data demonstrate a compartmentalization of injury processes. Innate immune cells including macrophages were located in glomerular and tubulointerstitial compartments, whereas, adaptive immune cells including memory B cells localized to the tubulointerstitial compartment. Furthermore, podocyte transcripts were decreased in glomeruli and did not recover with treatment indicating a vulnerability of these cells to acute AMR.

## Introduction

Current diagnosis of rejection has been confined to semiquantitative descriptions of histological findings or molecular analyses of homogenized tissue biopsies ^1^. Although both approaches have provided valuable insights to the diagnosis and potential mechanisms of different types of rejection, each has critical shortcomings. Histological analysis allows localization of different aspects of tissue injury and response, but limited insight about mediators. In contrast, molecular analysis of tissue homogenates disrupts pertinent evidence regarding localization of the molecular signals. Digital spatial profiling (DSP) bridges the advantages of these two approaches by resolving transcript analysis to localized histologically evaluable structures ^2^. Spatial profiling is particularly relevant in the kidney because of its intertwined compartments comprised of specialized cells. Spatial imaging combined with single-cell and single-nucleus assays has resolved 51 main cell types in human kidneys that can be altered by various disease processes ^3^. Heterogeneity in the distribution of pathological features has been recognized in the diagnosis of antibody mediated rejection (AMR). For example, a Banff g2 lesion score for glomerulitis requires *“*complete or partial occlusion of 1 or more glomerular capillary by leukocyte infiltration and endothelial cell enlargement” in 25 to 75% of glomeruli which can be segmental or global ^4^. This raises the question of differences in gene expression between glomeruli that do and do not display these histopathological features within a single biopsy. Therefore, we have applied digital spatial profiling to localize molecular signatures of cell injury to individual glomerular or tubular sites in sequential biopsies from a single patient before, during AMR and after treatment for AMR. Most DSP studies to date have been cross-sectional and used unrelated biopsies as controls. In this study, longitudinal biopsies from the same patient provided controls for the rejection analysis.

## Materials and Methods

### Serial biopsies from single patient

A series of 4 biopsies were available from our index patient (details in supplementary Table 1). A baseline biopsy from 25 days after transplantation followed by a 3 month biopsy diagnosed as acute AMR that was characterized by strong C4d staining of the glomeruli and moderate staining of the peritubular capillaries. After treatment with IVIg, a followup biopsy 4 months later revealed ongoing glomerulitis with a mild increase in tubulointerstitial fibrosis. A fourth biopsy 7 months later demonstrated resolved glomerulitis.

### GeoMx-Digital Spatial Profiling

Digital Spatial Profiling was performed according to protocols established by GeoMx (NanoString Inc, Seattle, WA) on excess formalin fixed paraffin embedded tissue (see supplementary detailed methods). The institutional review board of Cleveland Clinic approved the protocol. After hybridizing RNA Probes, immunofluorescent antibodies to the following targets were used to define regions of interest (ROIs): CD31(JC/70A abcam, Waltham, MA), CD68 (KP, Santa Cruz Biotechnology, Dallas, TX), and alpha smooth muscle actin (SP171 abcam, Waltham, MA). Each ROI was exposed to UV light to cleave the oligo associated barcods. Whole exome sequencing for 18,000+ protein-coding genes was performed using the GeoMX Digital Space Profiling platform (NanoString, Seattle, WA). GeoMx Data Analysis suite software (Nanostring; Version 3.0.0.113) was used to analyze the data.

Box and whisker visualization was created using Python with the Pandas, Matplotlib, and Seaborn libraries. Statistical analyses were conducted using non-parametric tests due to the non-normal distribution of data.

## Results

### Glomerular transcripts increased during rejection

Sections from the formalin fixed biopsies were stained initially with antibodies to CD31 and CD68 to select ROIs that were drawn around whole glomeruli (Fig 1A).

**Figure 1.**
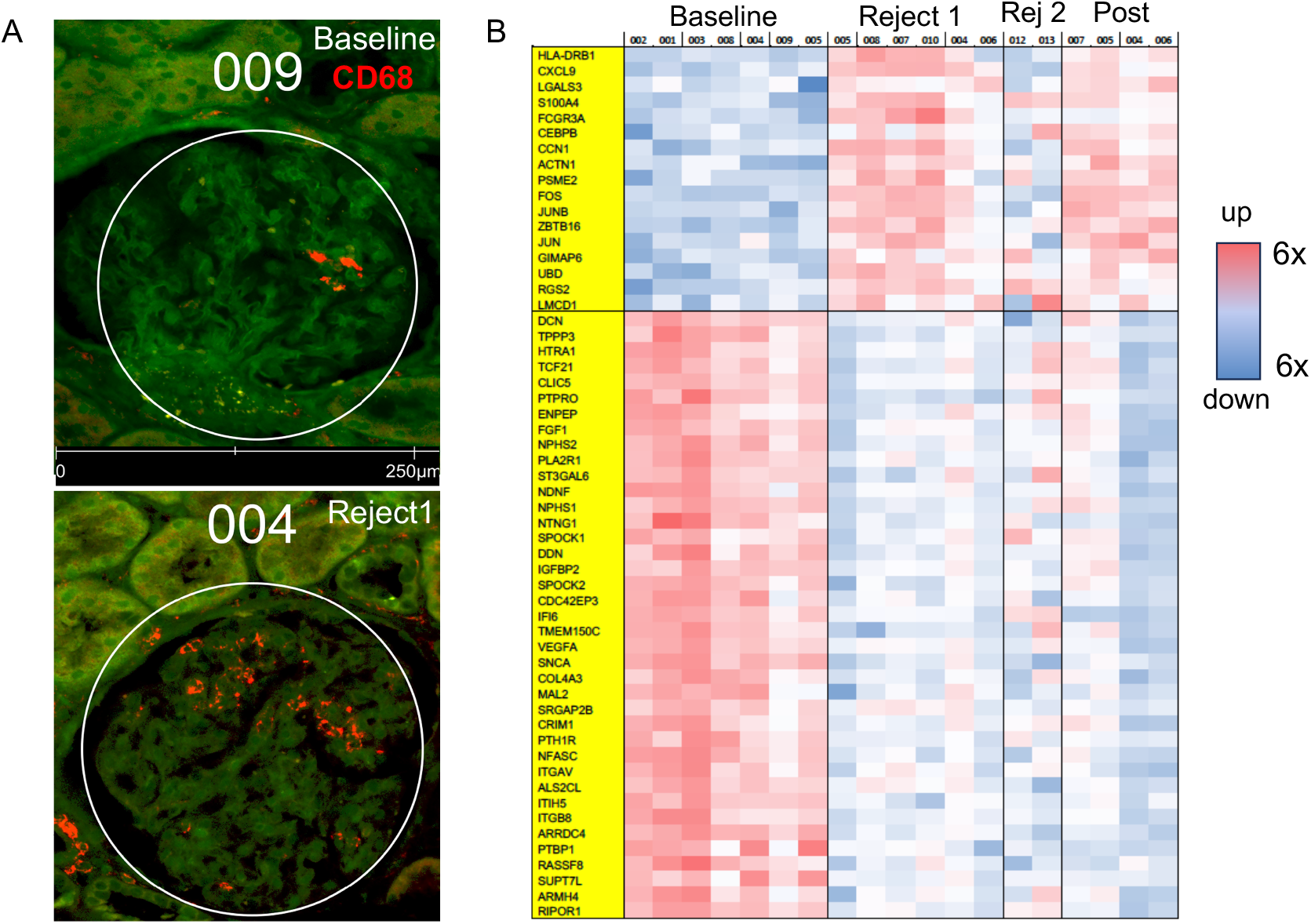
Glomerular transcripts in sequential biopsies: Baseline, AMR rejection (Reject 1), unresolved rejection (rej 2), Recovery (Post). **(a)** Representative glomeruli from baseline and Reject 1 containing CD68 positive (red) cells. **(b)** Heatmap of transcripts that differed from baseline by > 2-fold and p< 0.005 including 17 genes that increased from baseline (top) and 39 that decreased (bottom). Each column represents data from an individual glomerulus.

The baseline and 3 month biopsy (Rejection 1) provided 7 and 6 glomeruli, respectively, that passed quality control for reliable transcript analysis. Data were obtained from 2 and 4 glomeruli from the subsequent biopsies (ongoing rejection and resolution). Using GeoMx software, genes that differed from the baseline biopsy by > 2-fold with a significance of p<0.005 were identified. This resulted in 2 distinct sets of genes: 17 genes that increased from baseline and 39 that decreased (Fig 1B).

Deconvoluting the cell content of the ROIs using GeoMx software indicated that innate immune cells increased in glomeruli during the initial AMR episode. Increased signatures for macrophages (p=0.0175) correlated with the increased numbers of CD68 positive cells imaged in the corresponding ROIs (Fig 2A and B; Suppl Fig 1).

**Figure 2:**
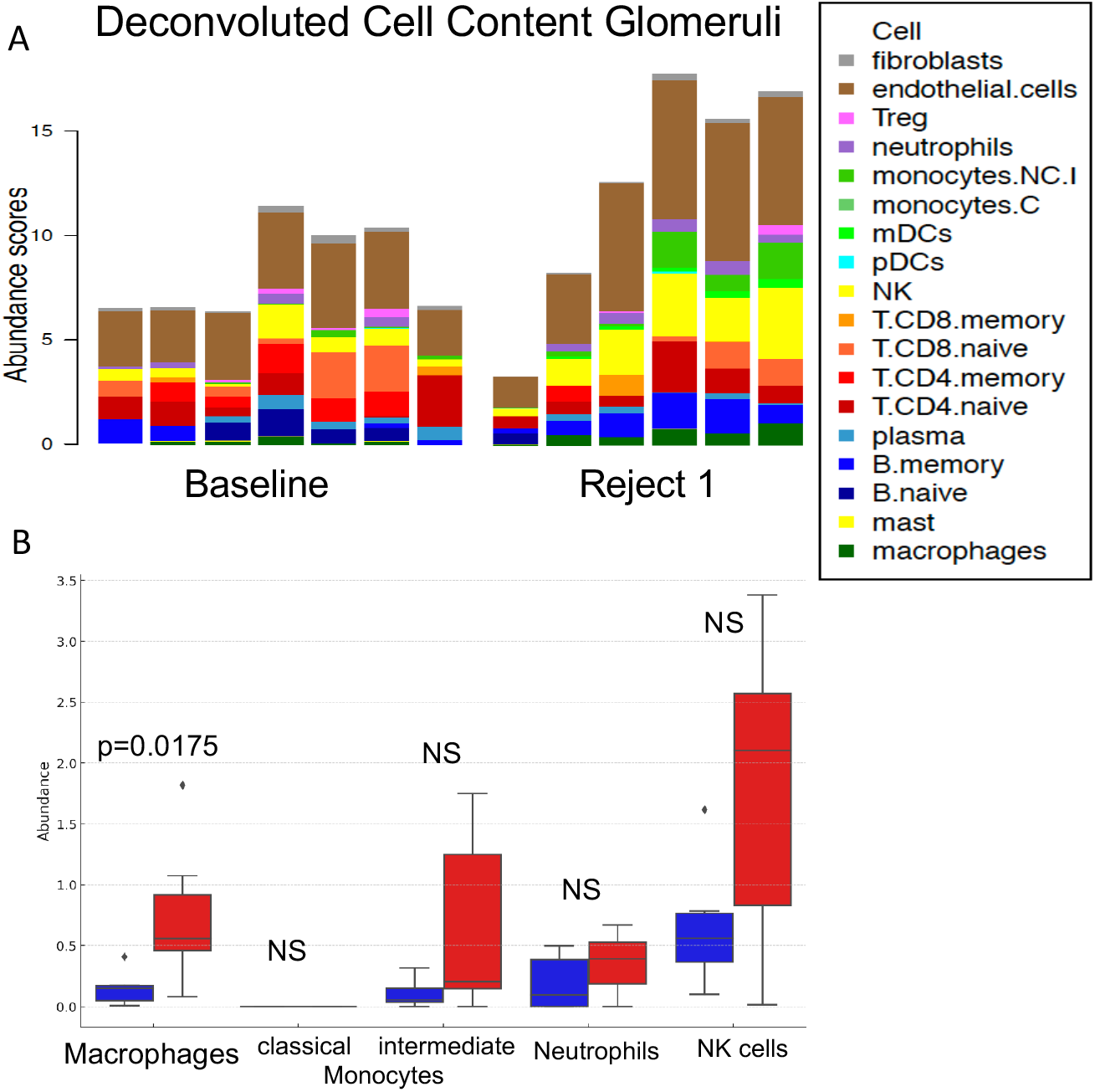
Deconvoluted cell content of Glomerular ROIs. **(a)** Stacked bar graphs for individual glomeruli show increased signatures for NK cells (yellow) as well as monocytes (green) and macrophages (dark green) in glomeruli during AMR (Reject1). **(b)** Box and whisker plots of abundance scores for macrophages, monocytes, neutrophils and NK cells created using Python with the Pandas, Matplotlib, and Seaborn libraries. Statistical analyses were conducted using non-parametric tests due to the non-normal distribution of data. Blue boxes = Baseline; Red boxes = Reject 1.

NK cells and intermediate monocytes were increased in 4 of 6 glomeruli in the AMR biopsy, but the differences did not reach statistical significance for the whole group because of the wide range of values (Fig. 2B).

HLA-DRB1 was among the 17 transcripts upregulated in the glomeruli during rejection. More detailed analysis revealed that transcripts for HLA-B and DR increased in the glomeruli from the baseline biopsy to the initial AMR at p <0.01, while HLA-DP and DQ increased at p <0.05 (Suppl Fig 2). HLA-B is of note because antibodies to HLA-B55 and B8 increased when AMR was diagnosed (Suppl Table 1).

Of the 15 transcripts identified in the Banff Human Organ Transplant (B-HOT) panel for AMR that is derived from molecular analyses of homogenized kidney biopsies, only CXCL9 differed from the baseline biopsy by > 2-fold with a significance of p<0.005 (Fig. 1A). When the B-HOT panel for AMR versus no rejection ^5^ was specifically interrogated most of the 15 genes tended to increase from baseline to AMR but at a lower level and only 7 reached a significance of p<0.05 (Suppl Fig 3).

### Glomerular transcripts decreased during rejection

Using the Human Cell Atlas, the 39 genes that were decreased during the initial rejection were determined to be expressed by podocytes, with many being signature genes for podocytes, such as NPHS1 (nephrin) and NPHS2 (podocin). Two other genes that have been identified as curated podocyte markers (NTNG1 and CLIC5) were also decreased (Fig 3). Analyzing the cell content of the ROIs by Python with the Pandas, Matplotlib, and Seaborn libraries corroborated the decrease in podocytes during acute AMR compared to the protocol biopsy (Fig 3B). Expression of these genes did not recover in the subsequent 2 biopsies.

**Figure 3.**
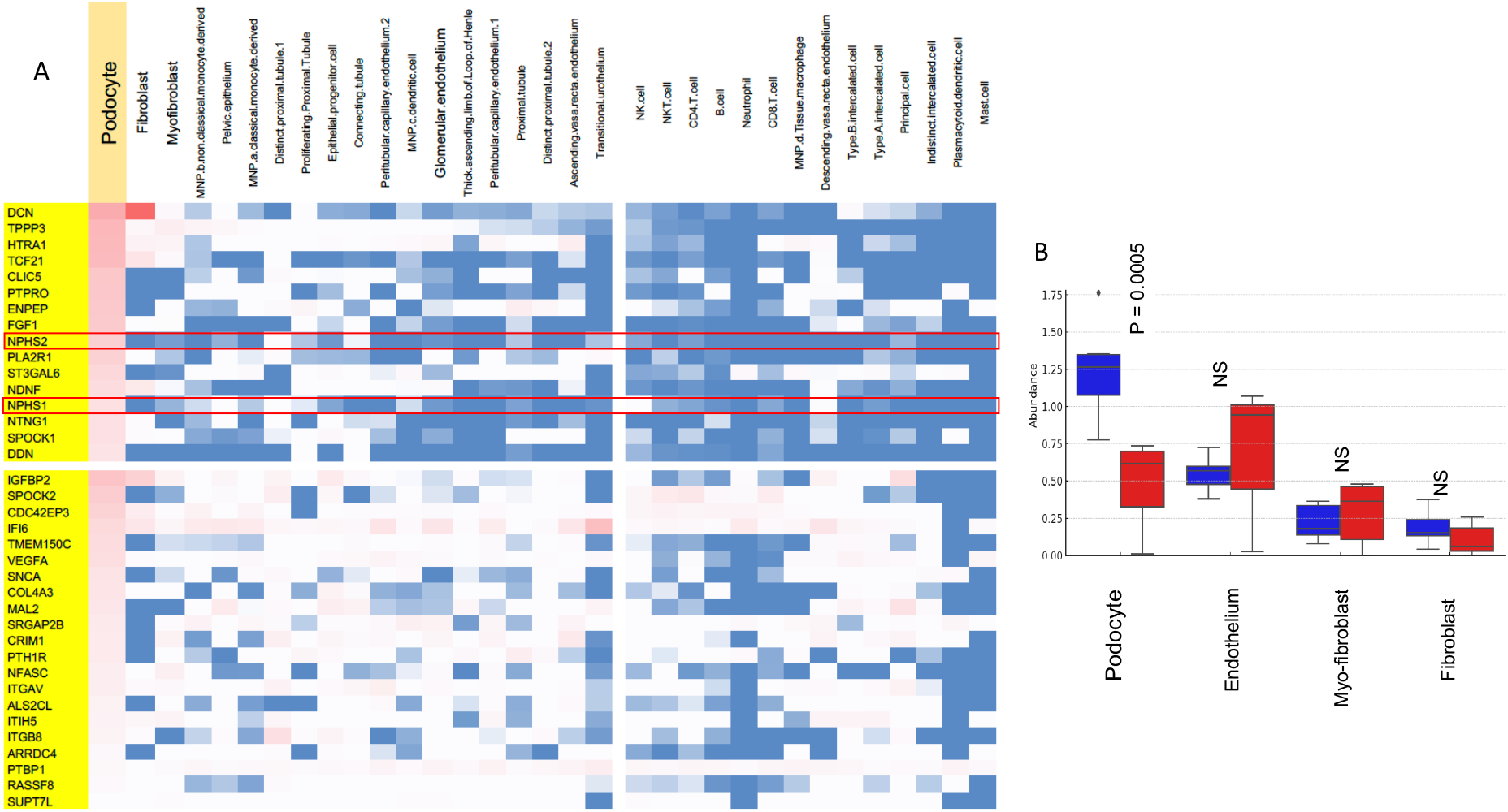
Glomerular transcripts downregulated in AMR rejection are associated with podocytes. **(a)** Heatmap for transcripts associated with different types of renal cells generated from the human cell atlas (HCA) indicate that 16 of the genes are predominantly expressed by podocytes (top). NPHS1 (nephrin) and NPHS2 (podocin) are signature genes for podocytes. Although the remaining genes are more widely expressed, they are also expressed by podocytes (bottom). **(b)** Box and whisker plots of abundance scores. Blue boxes = Baseline; Red boxes = Reject 1.

### Transcripts increased in areas of tubulointerstitial fibrosis during rejection

Additional ROIs encompassing tubulointerstitial fibrosis were defined using a marker antibody for alpha smooth muscle actin (Fig 4A). Successful segmenting was confirmed by the high expression of ACTA2 (SMA gene) and 3 additional genes characteristic of myofibroblasts and fibroblasts (TAGLN, NDUFA4L2 and NOTCH3). Deconvoluting the cell content of the segments with tubulointerstitial fibrosis disclosed significant signatures for memory B cells in the initial AMR sample. Treatment with IVIg did not eliminate the signal for memory B cells in the subsequent biopsy (Rejection 2 in Fig 4B). Signatures for macrophages and memory CD8 T cells also were increased in the fibrotic areas during ongoing rejection.

**Figure 4.**
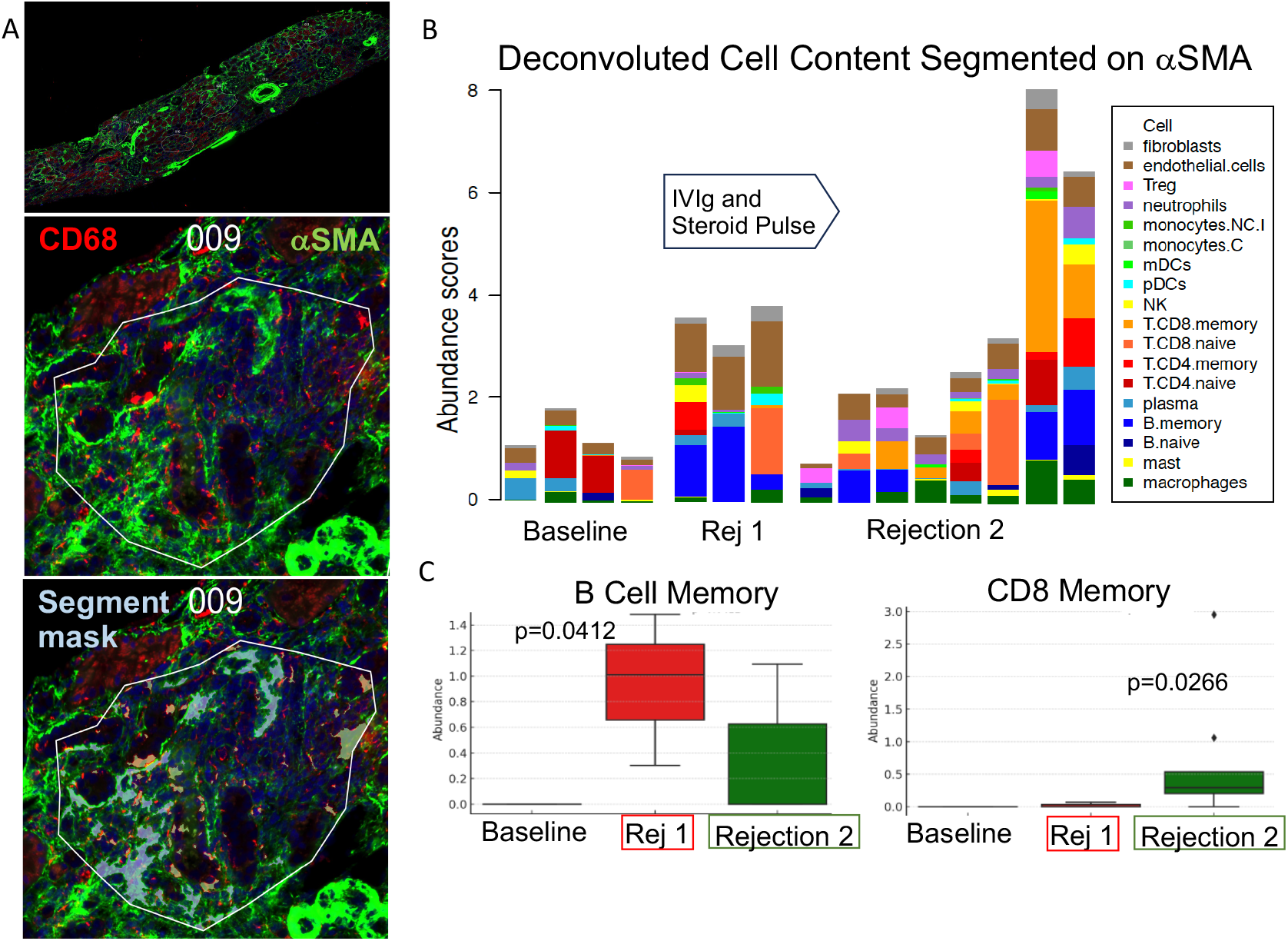
Transcripts increased in areas of tubulointerstitial fibrosis during rejection. **(a)** Tubulointerstitial fibrosis defined by alpha smooth muscle action (αSMA; green) in representative low power image of renal biopsy (top panel). Middle panel illustrates representative ROI encompassing tubulointerstitial fibrosis defined using a marker antibody for αSMA (green). ROI with mask (gray-blue) indicating segmentation on αSMA (Lower panel) **(b)** Deconvoluted cell content of the segments with tubulointerstitial fibrosis disclosed signatures for memory B cells (dark blue) in the initial AMR sample (Rej1) that remained after treatment with IVIg and pulse steroids (Rejection 2). Signatures for memory CD8 T cells (orange) and macrophages (dark green) also were increased in the fibrotic areas during ongoing rejection (Rejection 2). **(c)** Box and whisker plots of abundance scores for B and CD8 memory cells using Python with the Pandas, Matplotlib, and Seaborn libraries; Statistical analyses were conducted using non-parametric tests due to the non-normal distribution of data.

## Discussion

The kidney is a complex organ with multiple compartments that can be differentially involved by pathological processes ^3^. Moreover, the pathology is often not diffuse within a compartment but focal. The Banff criteria for renal allograft pathology recognize the significance of these different compartments by scoring the glomeruli, interstitium, tubules and vessels separately. The focal or segmental distribution is further scored by the percentage of each compartment that is injured. However, immunohistochemistry has limitations in the scope of probes available to assess pathological processes on a molecular level. Incorporation of microarray assessment of RNA extracted from renal biopsies resulted in a B-HOT panel expression of 770 genes which have been refined to correlate with different types of rejection ^6^. However, these molecular signatures cannot be assigned to specific structures within the kidney. In the current study, we were able to compare the molecular changes in glomerular and tubulointerstitial compartments from a baseline biopsy through acute AMR, treatment and resolution.

Spatial resolution of the transcriptome demonstrated that molecular signatures of innate immune cells including NK cells, monocytes and macrophages are located in glomerular compartments. These infiltrates were linked with a significant decrease in signatures for podocytes. The podocyte transcripts remained low even after treatment. This coincided with intense C4d staining in the glomeruli during AMR. Preponderant glomerular staining by C4d has been noted by Nankivell and colleagues in AMR as early as 1 month and extending through the first year after transplantation ^7^. The vulnerability of podocytes to AMR has been noted by Wiggins and colleagues ^8^.

Although transcripts associated with NK cells have been linked to AMR ^9^, it is noteworthy that transcripts for NK cells tended to increase in the glomerular but not tubulointerstitial compartments. Moreover, the signal for NK cells was heterogeneous with very high signal in 4 glomeruli and lower signal in 2 glomeruli. A similar but less pronounced variation was found for intermediate monocytes.

Significant differences were also found in the distribution of adaptive immune cells. Transcripts for memory B and CD8 T cells localized to the tubulointerstitial compartment amid myofibroblasts. A similar increase in B and T cells has recently been reported in areas of interstitial fibrosis using single-nuclei RNA-sequencing on renal biopsies ^10^. In our study, DSP demonstrated that treatment of the initial episode of AMR with pulse steroids and IVIg did not eliminate the signal for memory B cells in the subsequent biopsy. In addition, memory T cells expanded after treatment. In cell-mediated rejection expanded clones of T cells have also been found to resist therapeutic treatments ^11^.

Although this study is limited to a single patient, the availability of serial samples starting with an early protocol biopsy containing no evidence of rejection provided an ideal baseline to evaluate changes associated with AMR. The additional biopsies containing early tubulointerstitial fibrosis following treatment provided insights to the transition from acute AMR to chronic changes. Importantly, this case provided a spatial context for previous microarray findings.

## Supporting information

Supplemental text detailed methods

## DATA AND CODE AVAILABILITY

The data will be deposited in National Center for Biotechnology Information’s Gene Expression Omnibus.

## Author Contributions

Design of research study: RLF & WMB

Evaluating Patient samples and data: LCH, JJA

Conducting Digital Spatial Profiling: DM, HN & WMB

Analyzing data: DM, SK, ART, RLF & WMB

Writing the manuscript: PSH, ART, RLF & WMB

## ACKNOWLEDGMENTS

This work was supported by the CTOT GeoMx Digital Spatial Profiling Core (UO1 AI063594 to PSH and RLF) and grants NIH R01 AI165513 (WB) R01 AI158421 and AI40459 (RLF) from the NIAID of the NIH.

## References

1. Mengel M, Loupy A, Haas M, et al. Banff 2019 Meeting Report: Molecular diagnostics in solid organ transplantation-Consensus for the Banff Human Organ Transplant (B-HOT) gene panel and open source multicenter validation. Am J Transplant 2020; 20: 2305–2317.

2. Salem F, Perin L, Sedrakyan S, et al. The spatially resolved transcriptional profile of acute T cell-mediated rejection in a kidney allograft. Kidney Int 2022; 101: 131–136.

3. Lake BB, Menon R, Winfree S, et al. An atlas of healthy and injured cell states and niches in the human kidney. Nature 2023; 619: 585–594.

4. Haas M, Sis B, Racusen LC, et al. Banff 2013 meeting report: inclusion of c4d-negative antibody-mediated rejection and antibody-associated arterial lesions. Am J Transplant 2014; 14: 272–283.

5. Varol H, Ernst A, Cristoferi I, et al. Feasibility and Potential of Transcriptomic Analysis Using the NanoString nCounter Technology to Aid the Classification of Rejection in Kidney Transplant Biopsies. Transplantation 2023; 107: 903–912.

6. Smith RN, Rosales IA, Tomaszewski KT, et al. Utility of Banff Human Organ Transplant Gene Panel in Human Kidney Transplant Biopsies. Transplantation 2023; 107: 1188–1199.

7. Nankivell BJ, P’Ng CH, Shingde M. Glomerular C4d Immunoperoxidase in Chronic Antibody-Mediated Rejection and Transplant Glomerulopathy. Kidney Int Rep 2022; 7: 1594–1607.

8. Yang Y, Hodgin JB, Afshinnia F, et al. The two kidney to one kidney transition and transplant glomerulopathy: a podocyte perspective. J Am Soc Nephrol 2015; 26: 1450–1465.

9. Hidalgo LG, Sis B, Sellares J, et al. NK cell transcripts and NK cells in kidney biopsies from patients with donor-specific antibodies: evidence for NK cell involvement in antibody-mediated rejection. Am J Transplant 2010; 10: 1812–1822.

10. McDaniels JM, Shetty AC, Kuscu C, et al. Single nuclei transcriptomics delineates complex immune and kidney cell interactions contributing to kidney allograft fibrosis. Kidney Int 2023; 103: 1077–1092.

11. Shi T, Burg AR, Caldwell JT, et al. Single-cell transcriptomic analysis of renal allograft rejection reveals insights into intragraft TCR clonality. J Clin Invest 2023; 133.

