## Supplemental Table 1 for "Compartmentalization of Transcripts During Antibody Mediated Rejection in Renal Transplants"

**Supplementary Table 1:** Detailed Patient History:

Second kidney transplant to male recipient: Deceased donor, PRA 100%, Low levels of DSA to B55 and B8 pretransplantation but negative flow cytometry crossmatch

| Biopsy | Time | Diagnosis | DSA | MFI^a^ | Banff Scores | Treatment |
| --- | --- | --- | --- | --- | --- | --- |
| No biopsy | Pre-Transplt |  | B55  B8 | 1287  1959 |  |  |
| Baseline | 25 days | No Rejection | B55  B8  Cw9  DPB1*04:01 | 6675  8233  4890  1493 | i0, t0, v0,  g0, ptc0, c4d0,  ci0, ct0, cv0, cg0, ah0 | maintenance immunosuppression:  Tacrolimus  MMF  prednisone |
| Reject 1 | 3 months | Acute AMR | B55  B8  Cw9  DPB1*04:01  DQ5^b^ | 13415  17127  14639  1969  4020 | i0, t0, v0,  **g2, ptc1, c4d3**,  ci0, ct0, cv0, cg0, ah0 | Tacrolimus  MMF  Prednisone  pulse dose steroids IVIG |
| No biopsy | 5 months |  | B55  B8  Cw9  DQ5 | 4295  3979  7756  1112 |  |  |
| Reject 2 | 7 months | AMR not fully resolved and development of mild fibrosis |  |  | i0, t0, v0,  **g2, ptc0, c4d0**,  **ci1, ct1**, cv0, cg0, ah0 | Tacrolimus  MMF  Prednisone |
| No Biopsy | 11 months |  | B55  B8  Cw9  DQ5 | 4100  4121  7600  neg |  |  |
| Post | 14 months | Resolution of glomerulitis |  |  | i0, t0, v0,  **g0, ptc0, c4d0**,  **ci1, ct1**, cv0, cg0,ah0 | Tacrolimus  MMF  Prednisone |

^a^ Mean Fluorescence Index

^b^(DQB1*05:01 DQA1*01:01)
