## Supplemental text detailed methods for "Compartmentalization of Transcripts During Antibody Mediated Rejection in Renal Transplants"

Digital Spatial Profiling was performed according to protocols established by GeoMx (NanoString Inc, Seattle, WA) on excess formalin fixed paraffin embedded tissue. The institutional review board of Cleveland Clinic approved the use of excess tissue. Slides were prepared according to the GeoMx DSP Slide Preparation Manual for RNA. After hybridizing RNA Probes to the tissues overnight at 37C, unbound probes were removed and the hybridized slides underwent antigen retrieval before being stained with the following immunofluorescent antibodies: alexa fluor 647 conjugated mouse monoclonal antibody to CD31(JC/70A abcam, Waltham, MA), alexa fluor 594 conjugated mouse monoclonal antibody to CD68 (KP, Santa Cruz Biotechnology, Dallas, TX), alexa fluor 647 rabbit monoclonal to alpha smooth muscle actin (SP171 abcam, Waltham, MA), and alexa fluor 488 DNA stain (Syto13, Nanostring, Seattle, WA) to guide the selection of regions of interest (ROIs). Each ROI was exposed to a UV LED light to cleave the associated barcoded oligos. Whole exome sequencing was performed using the GeoMX Digital Space Profiling platform (NanoString, Seattle, WA). GeoMx Data Analysis suite software (Nanostring; Version 3.0.0.113) was used to analyze the data.

Box and whisker visualization was created using Python with the Pandas, Matplotlib, and Seaborn libraries. Statistical analyses were conducted using non-parametric tests due to the non-normal distribution of data.
