## Supplemental Figure 1 for "Compartmentalization of Transcripts During Antibody Mediated Rejection in Renal Transplants"

Suppl Fig 1: The expression profile of the 17 genes upregulated during AMR in the Human Cell Atlas for kidney.

| | Podocyte | Fibroblast | Myofibroblast | Proliferating.Proximal.Tubule | Proximal.tubule | Thick.ascending.limb.of.Loop.of.Henle | Indistinct.intercalated.cell | Distinct.proximal.tubule.2 | Principal.cell | Type.A.intercalated.cell | Peritubular.capillary.endothelium.2 | Descending.vasa.recta.endothelium | Glomerular.endothelium | Peritubular.capillary.endothelium.1 | Pelvic.epithelium | Distinct.proximal.tubule.1 | Epithelial.progenitor.cell | Connecting.tubule | Type.B.intercalated.cell | Ascending.vasa.recta.endothelium |
|---|---|---|---|---|---|---|---|---|---|---|---|---|---|---|---|---|---|---|---|---|
| S100A4 | 4.4 | 4.7 | 6.0 | 0.2 | 0.2 | 0.9 | 0.5 | 0.6 | 4.7 | 0.3 | 7.1 | 5.5 | 3.3 | 1.1 | 5.8 | 0.3 | 3.0 | 1.4 | 1.2 | 3.9 |
| FCGR3A | 0.3 | 0.0 | 0.0 | 0.1 | 0.0 | 0.1 | 0.0 | 0.1 | 0.0 | 0.0 | 0.1 | 0.0 | 0.0 | 0.1 | 0.0 | 0.0 | 0.1 | 0.0 | 0.1 | 0.1 |
| HLA-DRB1 | 0.1 | 0.6 | 1.4 | 1.3 | 1.4 | 0.2 | 0.1 | 3.2 | 1.8 | 0.3 | 8.2 | 10.4 | 13.2 | 14.1 | 4.5 | 0.7 | 1.0 | 1.1 | 0.0 | 13.9 |
| JUNB | 11.1 | 11.1 | 8.8 | 0.2 | 0.5 | 0.9 | 0.8 | 2.1 | 3.4 | 2.9 | 5.7 | 6.7 | 7.6 | 6.7 | 7.5 | 5.8 | 5.9 | 8.2 | 4.5 | 7.0 |
| FOS | 12.3 | 12.3 | 9.7 | 0.4 | 0.4 | 0.4 | 0.5 | 2.8 | 3.5 | 4.0 | 4.9 | 5.2 | 5.4 | 5.7 | 6.3 | 6.5 | 7.0 | 10.8 | 7.3 | 7.3 |
| RGS2 | 0.3 | 0.4 | 0.5 | 0.0 | 0.1 | 0.1 | 0.0 | 0.1 | 0.3 | 0.2 | 0.5 | 0.6 | 0.3 | 0.4 | 0.1 | 0.7 | 0.2 | 0.0 | 0.4 | 0.0 |
| CEBPB | 2.1 | 3.6 | 4.1 | 0.5 | 0.3 | 0.3 | 0.1 | 0.7 | 0.2 | 0.1 | 1.6 | 1.6 | 5.0 | 2.3 | 2.1 | 1.7 | 0.6 | 0.5 | 0.3 | 1.7 |
| PSME2 | 1.5 | 1.5 | 1.2 | 1.9 | 2.1 | 2.5 | 3.6 | 2.7 | 1.8 | 2.0 | 1.0 | 1.7 | 2.2 | 2.3 | 4.4 | 1.7 | 1.3 | 1.0 | 1.4 | 2.1 |
| JUN | 10.4 | 9.0 | 8.4 | 0.4 | 0.5 | 2.0 | 0.6 | 3.3 | 2.6 | 5.4 | 6.3 | 6.3 | 9.7 | 9.0 | 5.5 | 6.0 | 5.8 | 9.1 | 8.5 | 8.4 |
| LGALS3 | 0.2 | 6.4 | 1.9 | 0.8 | 0.8 | 1.8 | 9.9 | 3.2 | 1.5 | 8.9 | 2.5 | 3.4 | 0.5 | 1.2 | 5.7 | 0.8 | 2.7 | 2.9 | 7.9 | 1.4 |
| ZBTB16 | 1.3 | 1.3 | 4.3 | 0.2 | 0.2 | 0.3 | 0.2 | 0.7 | 0.1 | 0.8 | 2.2 | 2.6 | 5.7 | 3.6 | 0.0 | 2.5 | 1.4 | 0.2 | 1.5 | 2.0 |
| GIMAP6 | 0.0 | 0.0 | 0.2 | 0.0 | 0.0 | 0.0 | 0.0 | 0.4 | 0.0 | 0.0 | 1.1 | 1.0 | 2.0 | 1.9 | 0.0 | 0.0 | 0.0 | 0.1 | 0.0 | 1.5 |
| ACTN1 | 0.2 | 1.7 | 3.2 | 0.1 | 0.1 | 0.2 | 0.2 | 0.3 | 0.6 | 0.1 | 0.5 | 0.4 | 0.1 | 0.1 | 0.5 | 1.6 | 0.7 | 0.2 | 0.2 | 1.4 |
| UBD | 0.0 | 0.0 | 0.0 | 0.0 | 0.0 | 0.0 | 0.0 | 0.2 | 0.0 | 0.0 | 0.0 | 0.0 | 0.0 | 0.0 | 0.4 | 0.4 | 0.2 | 0.0 | 0.0 | 0.0 |
| LMCD1 | 0.1 | 0.9 | 0.8 | 0.4 | 0.3 | 0.1 | 0.0 | 0.5 | 0.1 | 0.1 | 1.9 | 2.4 | 0.3 | 1.7 | 0.3 | 0.4 | 0.2 | 0.1 | 0.0 | 2.7 |
| CXCL9 | 0.0 | 0.0 | 0.0 | 0.0 | 0.0 | 0.0 | 0.0 | 0.0 | 0.0 | 0.0 | 0.0 | 0.0 | 0.0 | 0.0 | 0.0 | 0.0 | 0.0 | 0.0 | 0.0 | 0.0 |

| | Neutrophil | MNP.b.non.classical.monocyte.derived | MNP.a.classical.monocyte.derived | MNP.d.Tissue.macrophage | MNP.c.dendritic.cell | NKT.cell | NK.cell | CD8.T.cell | CD4.T.cell | Mast.cell | Transitional.urothelium | B.cell | Plasmacytoid.dendritic.cell |
|---|---|---|---|---|---|---|---|---|---|---|---|---|---|
| S100A4 | 25.5 | 15.9 | 19.2 | 9.4 | 8.1 | 13.1 | 8.4 | 9.3 | 11.4 | 12.3 | 4.0 | 2.4 | 2.5 |
| FCGR3A | 9.4 | 11.1 | 1.5 | 5.5 | 3.3 | 5.3 | 8.8 | 0.5 | 0.1 | 0.0 | 0.0 | 0.0 | 0.0 |
| HLA-DRB1 | 0.8 | 10.6 | 10.2 | 24.8 | 19.7 | 2.1 | 0.6 | 2.6 | 0.6 | 0.0 | 1.7 | 13.7 | 8.0 |
| JUNB | 17.2 | 9.2 | 9.2 | 8.0 | 7.8 | 8.2 | 7.8 | 13.2 | 12.7 | 14.2 | 12.8 | 14.3 | 6.5 |
| FOS | 41.3 | 9.2 | 13.0 | 11.3 | 10.8 | 8.5 | 8.2 | 8.5 | 9.9 | 16.7 | 12.9 | 7.4 | 7.3 |
| RGS2 | 38.1 | 8.5 | 10.0 | 8.4 | 8.5 | 3.7 | 4.5 | 3.5 | 3.4 | 18.7 | 0.0 | 4.3 | 9.7 |
| CEBPB | 10.3 | 7.2 | 6.7 | 4.3 | 2.8 | 2.1 | 2.0 | 1.3 | 1.2 | 1.0 | 0.8 | 0.7 | 0.0 |
| PSME2 | 0.0 | 5.1 | 3.6 | 2.3 | 2.5 | 2.4 | 3.0 | 2.7 | 2.7 | 0.2 | 1.1 | 1.6 | 1.8 |
| JUN | 0.0 | 4.6 | 2.7 | 6.2 | 7.0 | 11.5 | 9.2 | 16.3 | 15.9 | 21.7 | 12.9 | 17.2 | 2.6 |
| LGALS3 | 0.0 | 3.4 | 4.0 | 3.3 | 2.0 | 0.2 | 0.2 | 0.8 | 1.0 | 3.8 | 4.3 | 0.2 | 0.0 |
| ZBTB16 | 0.7 | 1.1 | 0.7 | 0.8 | 1.3 | 0.2 | 1.6 | 0.3 | 0.4 | 0.2 | 0.0 | 0.2 | 0.0 |
| GIMAP6 | 0.0 | 0.4 | 0.2 | 0.1 | 0.1 | 0.5 | 1.0 | 0.8 | 0.7 | 0.0 | 0.0 | 0.0 | 0.2 |
| ACTN1 | 1.9 | 0.2 | 0.6 | 0.2 | 0.5 | 0.1 | 0.1 | 0.1 | 0.2 | 0.0 | 0.7 | 0.0 | 0.0 |
| UBD | 0.0 | 0.0 | 0.0 | 0.0 | 0.0 | 0.0 | 0.0 | 0.0 | 0.0 | 0.0 | 0.0 | 0.0 | 0.0 |
| LMCD1 | 0.0 | 0.0 | 0.1 | 0.0 | 0.1 | 0.0 | 0.1 | 0.0 | 0.1 | 0.2 | 0.0 | 0.0 | 0.0 |
| CXCL9 | 0.0 | 0.0 | 0.0 | 0.0 | 0.3 | 0.0 | 0.0 | 0.0 | 0.0 | 0.0 | 0.0 | 0.0 | 0.0 |
