## Supplemental Figure 2 for "Compartmentalization of Transcripts During Antibody Mediated Rejection in Renal Transplants"

Suppl Fig 2

HLA increased in the glomeruli in AMR

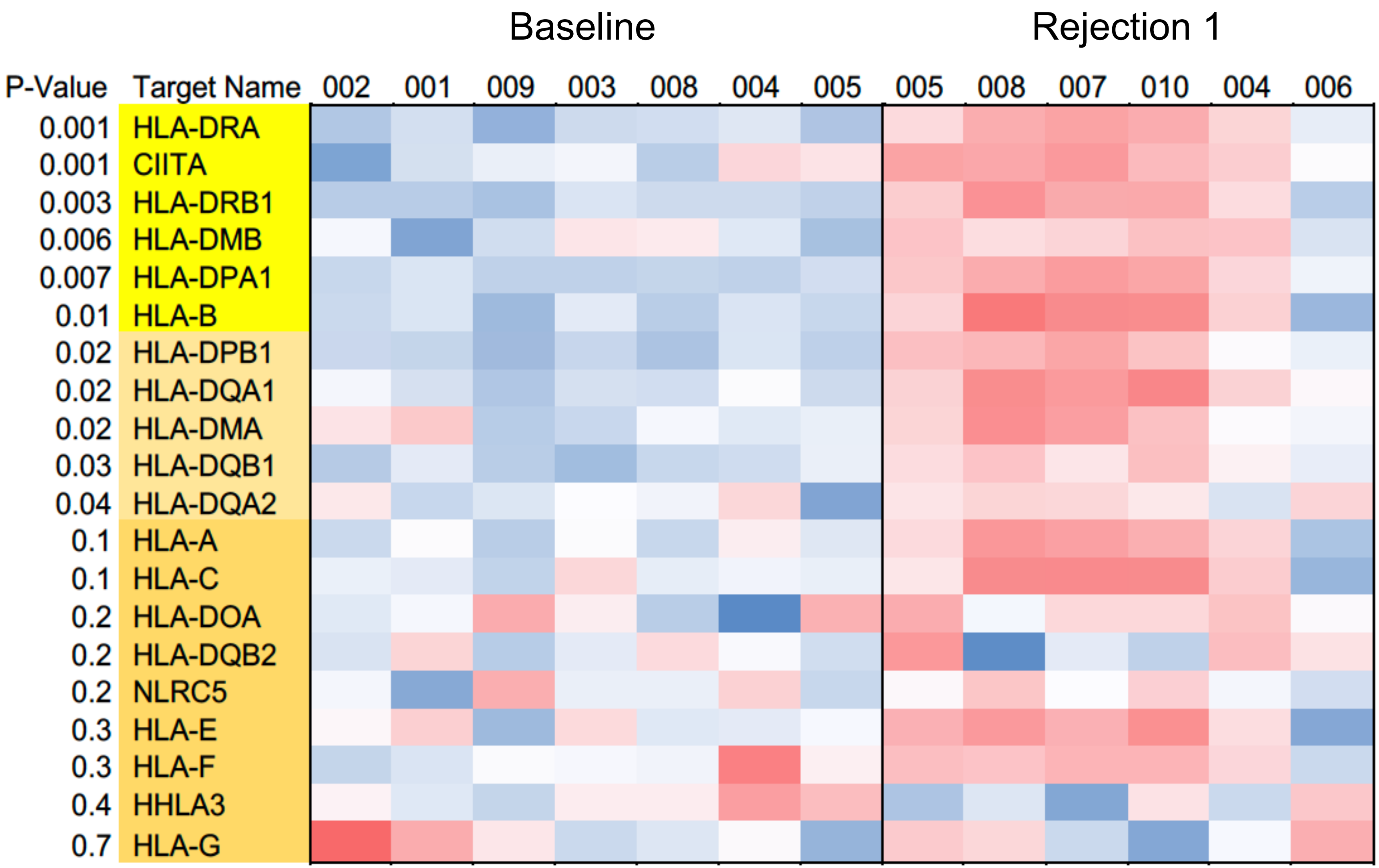


**Figure S2.** Heatmap of transcripts for HLA in glomeruli. HLA-B and DR transcripts reached a threshold of 2-fold and **p<0.01**; whereas HLA-DP and DQ reached a threshold of 2-fold and **p<0.05**.
