## Supplemental Figure 3 for "Compartmentalization of Transcripts During Antibody Mediated Rejection in Renal Transplants"

Suppl Fig 3

### B-HOT panel for AMR versus no rejection

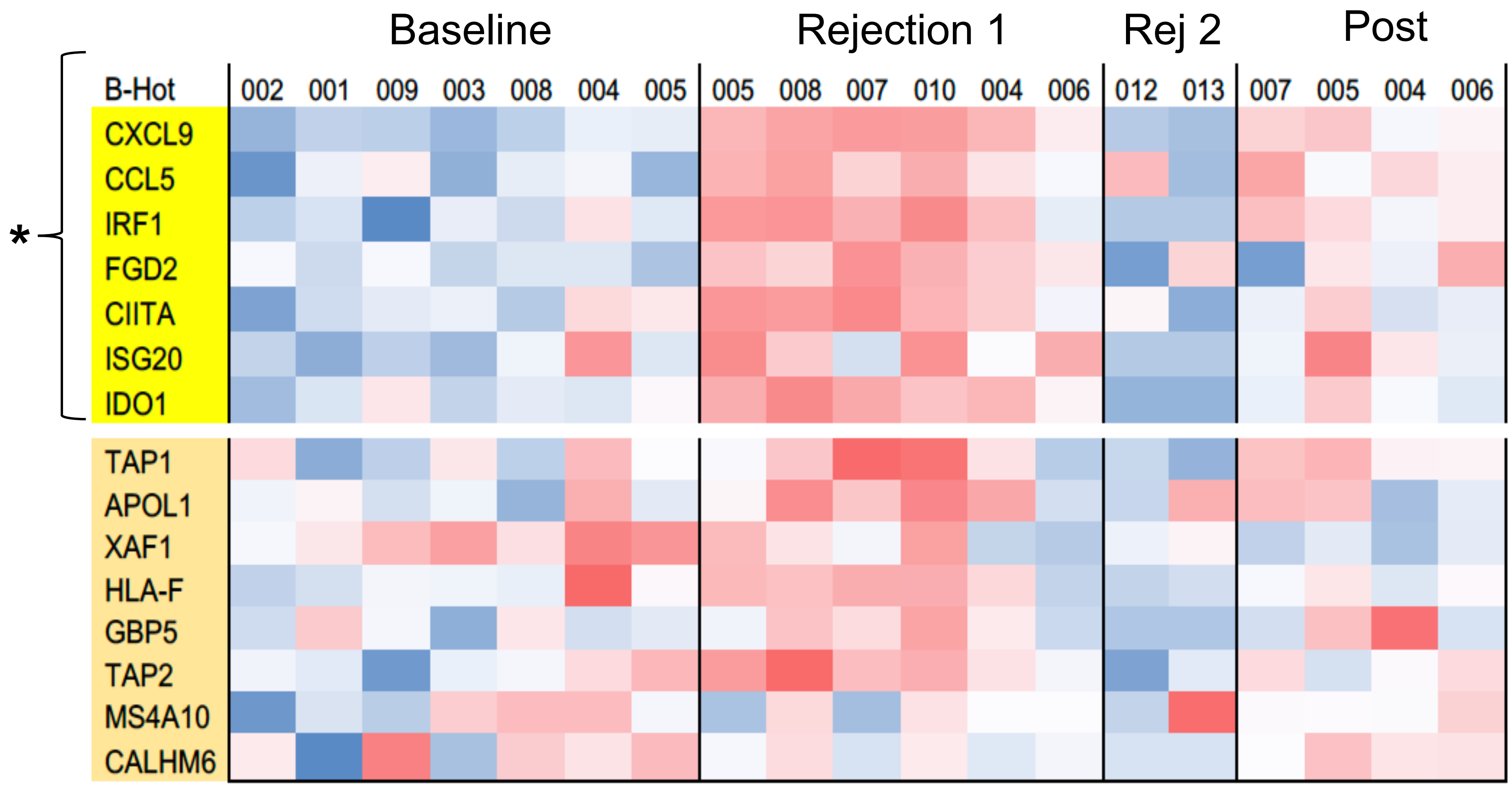


**Figure S3.** The 15 genes in the B-HOT panel for AMR versus no rejection tended to increase from baseline to AMR but at a lower level and only 7 reached a significance of $p<0.05$*.
